# High cumulative viral titers of influenza virus in animals with significant disease fatality rates indicate a potential trade-off between fatality and transmissibility

**DOI:** 10.1101/2024.12.11.627227

**Authors:** Ayrton S. Gouveia, Leonardo G. Mata, Daniel A.M. Villela

## Abstract

Evaluating the trade-off hypothesis for the evolution of virulence using empirical data poses significant challenges. The hypothesis suggests that pathogens evolve to maximize transmissibility, but fatality imposes limits as there are diminishing gains in transmissibility. In this study, we analyzed a secondary dataset of influenza virus infections in ferrets (*Mustela putorius furo*), categorized by Hemagglutinin (HA) and Neuraminidase (NA) subtypes. Subgroups defined by the H7/N9 and H7/N7 combinations exhibited fatality rates of approximately 30% and reached cumulative viral titers close to 7.5 (log_10_ titer/mL). These levels represent intermediate fatality rates, as the H5/N6 and H5/N1 subgroups had higher fatality rates but reached lower cumulative viral titers. Using cumulative viral titer as a proxy for potential secondary transmissions, the analysis suggests that intermediate fatality rates are associated with higher numbers of secondary transmissions. However, there are significant uncertainties in subgroups with lower or no fatalities. Additionally, subgroups without fatalities showed substantial variability in cumulative viral titers.

## Introduction

Virulence in infectious diseases refers to a pathogen’s capacity to cause harm to its host. The persistence of a pathogen relies on its ability to sustain successive generations, which is ultimately determined by its fitness. Various factors may exert evolutionary pressure on pathogen traits for instance favoring traits that enhance the transmissibility. However, there are counterintuitive theories regarding the evolution of virulence. For instance, conditions that balance transmissibility and virulence may favor intermediate levels of virulence as optimal, as proposed by the trade-off hypothesis (1).

The harm potential given by virulence depends on factors such as the microorganism’s replication rate and immune evasion mechanisms. Levin and Bull (1994) (2) argue that the evolution of virulence is linked to a trade-off between causing enough harm to facilitate transmission and avoiding rapid host death, which would limit spread. Based on this balance, pathogens can evolve to optimal levels of virulence. According to Ewald (1994) (1), pathogens transmitted by vectors, such as mosquitoes, or through contaminated water can evolve to become more virulent, as host immobility does not hinder transmission. On the other hand, pathogens that rely on direct contact between hosts may be less virulent, as they require greater mobility from infected individuals to ensure further spread. Frank (1996) (3) also suggests that the mode of transmission influences virulence. Understanding how virulence manifests in animal models, as highlighted by Read (1994) (4), is crucial since these models provide a controlled environment for studying pathogen-host interactions, enabling more effective interventions.

The ferret (*Mustela putorius furo*) is widely regarded as an excellent model organism for studying influenza virus infection due to its ability to replicate key clinical signs observed in infected humans. Unlike other small mammals such as mice or guinea pigs, ferrets can become productively infected with various strains of human, avian, and zoonotic influenza A viruses (IAV) without requiring prior adaptation to the host (5). This characteristic enables researchers to investigate both viral pathogenicity and transmission in a single model, making ferrets invaluable for public health risk assessment and the study of emerging infectious diseases. Additionally, the ferret model allows for the study of virus transmissibility through multiple modes, such as respiratory droplets, mimicking the transmission routes observed in human populations. These features have solidified the ferret’s position as a gold standard in influenza research, used extensively by global health organizations like the CDC and WHO for pandemic preparedness and influenza risk assessment (6,7).

In this study, we aim to verify whether the trade-off hypothesis applies to the virulence of IAV in ferrets. We used a secondary dataset of ferrets used as animal models to investigate the Influenza A virus. This dataset has been previously used to study the spatial and temporal dynamics of IAV replication and spread within ferret hosts, as well as to predict outcomes using machine learning on in vivo data. It provides extensive information on respiratory droplet transmission events among animals, longitudinal viral titers, and fatality data, enabling a thorough evaluation of transmissibility and fatality levels across different IAV groups.

## Methodology

### Study population

We use in this work a public dataset (8) of 728 young male ferrets contaminated with 126 different types of influenza avian virus (IAV). The inoculation process was made intranasally with high doses (10^5^–10^7^ infectious units) of IAVs. The inoculated IAVs can be classified by their pathogenic level i.e highly pathogenic, and origins, i.e avian, swine, and canine. After the infection, those animals were monitored daily to track sublethal effects of the inoculated IAV, like an increase in their normal body temperature, and weight loss. On the other hand, the viral titer was evaluated from nasal wash (NW) collected on alternate days after 1 day post-inoculation (p.i) for 613 ferrets, and after 2 for 115. Those NW from 444 ferrets had the viral titer estimated using embryonated hen’s eggs and 284 on MDCK cells. Those two substrates result in two and interchangeable units. Among the Hemagglutinin (HA) and Neuraminidase (NA) subgroups of these IAVs, the most common were H1/N1 and H5/N1, observed in 174 and 144 ferrets, respectively.

A total of 108 ferrets died during the experiments, 95 of which were inoculated with a highly pathogenic virus. The survival time of the ferrets that died ranged from 3 to 13 days, with 82 of them dying between 5 and 9 days post-inoculation. Additionally, the capacity of an infected ferret to transmit the virus to a naive one was investigated in 230 cases, of which 103 resulted in transmission and 127 did not.

### Inclusion and exclusion criteria

For the analysis, two exclusion criteria were applied to the original dataset. First, any ferrets without viral titer measurements taken on alternate days following 1 day post-infection (p.i.) were excluded to standardize evaluation metrics and minimize bias in the estimation of average and maximum titer levels. The second criterion concerned the unit used for titer estimation and the variety of HA and NA subgroups. Only cases where titers were estimated in EID were included, as this ensured consistency across all HA and NA subtypes observed. Consequently, data from 350 ferrets were utilized in the statistical analysis.

### Statistical analysis

All analyses were conducted by grouping the HA and NA subtypes of each IAV into subgroups that describe the protein envelope of each virus. The data were first used to calculate summary statistics for the sublethal effects and viral titer, including means, standard deviations, and percentage. For the viral titer, we also assessed its variation over time. We fitted generalized linear models (GLMs) to estimate both transmissibility and recovery rates across all IAV subgroups. Additionally, survival analysis was conducted to calculate the mortality rates specific to each subgroup.

A logistic regression was used to analyze the transmission rate, using the transmission event as the dependent variable. This model included an interaction term between the maximum viral titer and the IAV subgroups to assess how this relationship affects transmissibility. However, this analysis was limited to the 230 ferrets for which the transmission event information was available.

For the recovery rate, we assumed that a viral titer below the cutoff of 2 log□□ titer/mL indicated recovery. This threshold allowed us to classify the ferrets as recovered based on their viral titer levels. Subsequently, the number of days for the viral titer to reach this threshold was used as the dependent variable with the IAV subgroups as independent variables in a Poisson regression. The daily mortality rates for each subgroup were calculated as the inverse of the exponentiated model’s results.

The fatality rate is obtained for each IAV subgroup by the number of deaths divided by the total infected animals. To estimate the daily mortality rate, we first calculated the survival probabilities of ferrets infected with each IAV subgroup using Kaplan-Meier estimator. Next, we calculated a general daily mortality rate for each subgroup by summing the number of deaths and dividing by the total observed time within that subgroup. It is important to note that only ferrets infected with the subgroups H5/N1, H5/N6, H7/N7, H7/N8, and H7/N9 died, which limited our estimated rates to these subgroups.

To estimate the maximum viral titer for each subgroup, a linear regression was applied, with the cumulative maximum viral titer as the dependent variable and the IAV subgroup categories as independent variables. The predicted maximum viral titer for each HA/NA subgroup was obtained by adding the coefficients to the model intercept. The precision of the estimates was assessed using 95% confidence intervals. This method allows for a detailed analysis of the differences in viral titers, enabling a comparison of the maximum viral replication in ferrets inoculated with the different subgroups.

The cumulative viral titer was obtained for all IAV subgroups over the study time. A measure of individual cumulative viral tier was summed over all surviving days for each ferret: for an individual *i*, the individual cumulative viral titer *C*_*i*_ is given by *C*_*i*_ = ∑_*j*_ λ_*ij*_, where *j* covers days. We evaluate the cumulative viral titer per subgroup by the geometric mean and reporting on logarithmic value 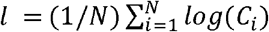, where N is the number of animals in the subgroup We evaluate the cumulative viral titer as a proxy of the reproduction number. For a ferret *f*_*i*_ with survival from day 1 to day *J*, the number of infected individuals on any given day *j* will be a function *g(*λ_*ij*_), where λ_*ij*_ is the viral titer. In field scenarios, this function typically depends on the number of contacts of individuals and the probability of transmission upon contact. Assuming the number of animals infected from a single ferret to be proportional to its individual cumulative viral titer, the total infected individuals is 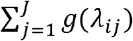 proportional to *C*_*i*_.

## Results

A total of 350 individuals were analyzed, distributed across 15 different subgroups defined by hemagglutinin and neuraminidase subtypes (Table 1). The percentage of respiratory droplet transmission varied significantly among the subgroups analyzed. In the H1/N1 subgroup, 7.5% of ferrets exhibited droplet transmission, while in the H3/N2, this proportion reached 47.37%. Some subgroups, such as H2/N3, showed minimal transmission (8.33%), while others recorded no transmission events. The mean temperature increase ranged from 0.73°C (± 0.31) in the H7/N1 subgroup to 2.34°C (± 0.95) in the H7/N3. Weight loss also showed significant variation, with the lowest value recorded being 2.20% (± 2.84) in the H7/N1 subgroup and the highest at 13.10% (± 8.45) in the H5/N6.

**Table 1.** Descriptive statistics of transmission events, symptoms, viral titer and deaths.

| HA/NA* | N | Transmission by<br>respiratory<br>droplets N (%) | Number of<br>individuals<br>without data<br>on<br>transmission<br>by respiratory<br>droplets N<br>(%) | Mean<br>temperature<br>(St.Dev.) ** | Mean weight<br>loss<br>(St.Dev.)<br>*** | Mean titer<br>(Sd.Dev.) | Deaths<br>(%) | Mean<br>survival<br>(Sd.Dev.) |
| --- | --- | --- | --- | --- | --- | --- | --- | --- |
| H1/N1 | 40 | 3 (7.5) | 19 (47.5) | 1.29 (0.95) | 6.18 (4.53) | 4.42 (2.05) | 0 | - |
| H2/N2 | 15 | 3 (20) | 10 (80) | 1.44 (0.79) | 4.26 (3.87) | 4.56 (2.57) | 0 | - |
| H2/N3 | 12 | 1 (8.33) | 6 (50) | 1.64 (0.54) | 7.65 (7.66) | 4.80 (2.20) | 0 | - |
| H3/N2 | 19 | 9 (47.37) | 10 (52.63) | 1.51 (0.88) | 5.89 (3.64) | 4.37 (2.15) | 0 | - |
| H5/N1 | 120 | 0 (0) | 108 (90) | 1.94 (0.73) | 12.80 (7.30) | 4.29 (1.83) | 41.67 | 6.72 (1.46) |
| H5/N2 | 6 | 0 (0) | 6 (100) | 1.35 (0.43) | 3.30 (5.06) | 3.39 (1.62) | 0 | - |
| H5/N6 | 9 | 0 (0) | 9 (100) | 2.07 (0.55) | 13.10 (4.15) | 4.08 (1.71) | 55.56 | 6.20 (2.77) |
| H5/N8 | 3 | 0 (0) | 3 (100) | 1.27 (0.31) | 3.53 (5.12) | 2.83 (1.62) | 0 | - |
| H7/N1 | 3 | 0 (0) | 3 (100) | 0.73 (0.35) | 2.20 (2.84) | 4.61 (1.94) | 0 | - |
| H7/N2 | 33 | 0 (0) | 21 (63.64) | 1.10 (0.54) | 6.48 (5.69) | 4.58 (2.23) | 0 | - |
| H7/N3 | 12 | 0 (0) | 9 (75) | 2.34 (0.93) | 12.80 (8.45) | 4.83 (2.11) | 0 | - |
| H7/N7 | 24 | 0 (0) | 18 (75) | 1.58 (0.76) | 12.60 (6.78) | 5.21 (2.02) | 29.17 | 7.57 (2.15) |
| H7/N8 | 6 | 0 (0) | 6 (100) | 1.38 (0.45) | 8.58 (9.21) | 3.82 (1.79) | 16.67 | 10.00 (-) |
| H7/N9 | 27 | 1 (3.7) | 18 (66.67) | 1.63 (0.67) | 11.30 (8.07) | 4.87 (1.96) | 33.33 | 8.78 (1.99) |
| H9/N2 | 21 | 7 (33.33) | 9 (42.86) | 1.64 (0.45) | 7.68 (4.32) | 5.58 (2.15) | 0 | - |

\*Hemagglutinin/Neuraminidase subgroups; \*\* increase on the ferrets standard body temperature; \*\*\* reduction on the ferrets standard weight. A dash means “not available”.

In terms of titration, the mean values ranged from 2.83 (± 1.62) in the H5/N8 subgroup to 5.58 (± 2.15) in the H9/N2, indicating different immune responses across subgroups.

Mortality rates also varied considerably. The H5/N6, H7/N9, H5/N1, and H7/N7 subgroups had the highest rates, at 55.56%, 33.33%, 41.67%, and 29.17%, respectively, while other subgroups did not record any deaths.

The mean survival period was reported for some subgroups with fatalities, with the longest survival observed in the H7/N8 subgroup at 10 days, followed by H7/N9 at 8.78 days, and H7/N7 at 7.57 days. These findings suggest significant variation in the infection response and progression across the different subtypes analyzed.

Figure 1 presents the viral titers (log10 titer/mL) of the 15 different HA/NA subgroups over nine days. Most subgroups reached peak viral titers between days 1 and 5 post-inoculation, followed by a gradual reduction until day 9.

**Figure 1.**
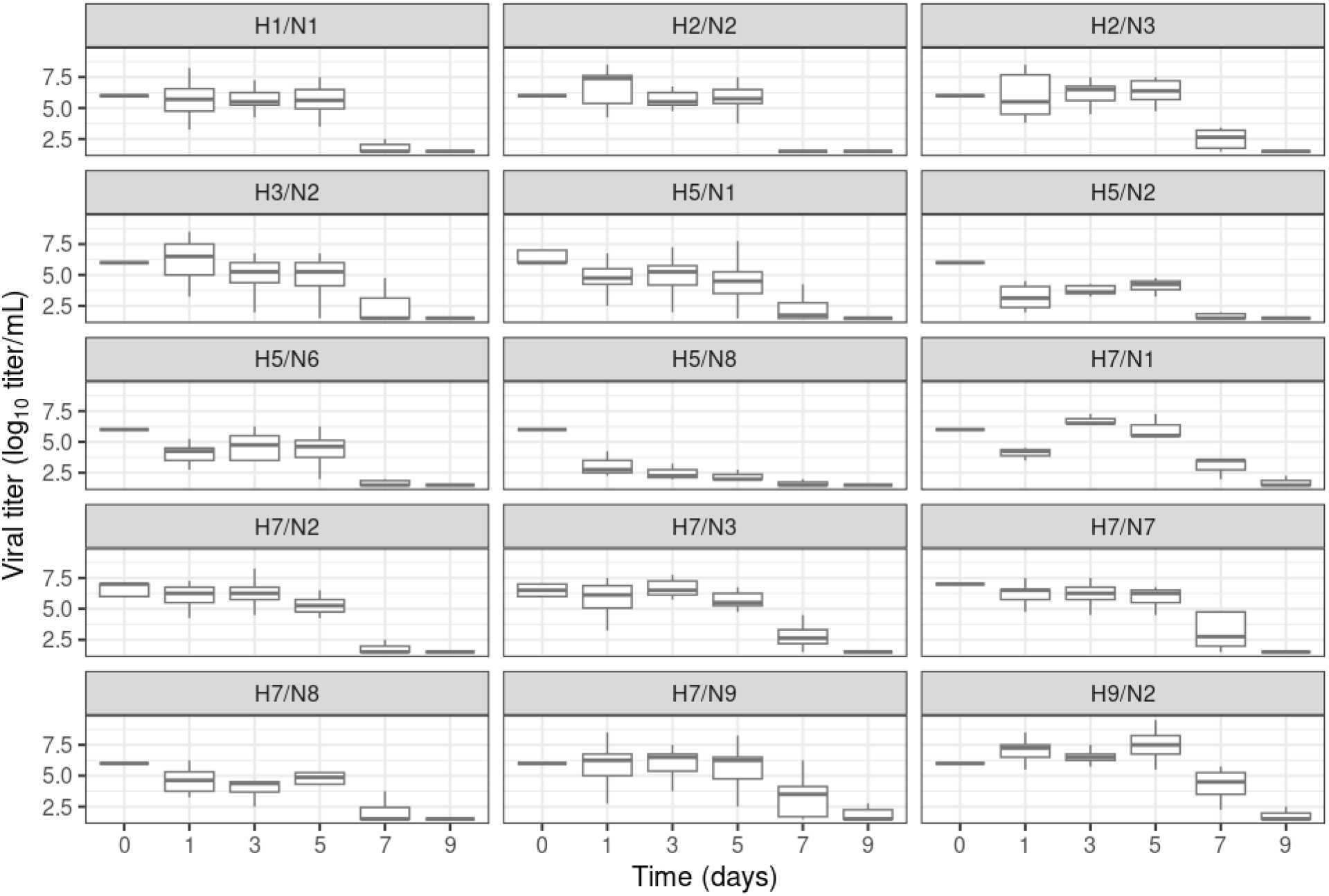
Distribution of viral titers by days post-inoculation separated by IAV subgroups.

The subgroups H5/N8, H7/N1, and H9/N2 stood out for their distinct behavior. The H5/N8 subgroup showed a decrease in viral titers starting on day 1, maintaining this downward trend until day 9. The H7/N1 subgroup showed an initial decrease on day 1, followed by an increase on day 3, and then a decline again on day 5, with a gradual reduction continuing until day 9. On the other hand, the H9/N2 subgroup exhibited a more stable pattern in the early days, with a rapid increase on day 5, followed by a sharp reduction on day 7, continuing to decline through day 9.

These results suggest that, while most subgroups follow a similar pattern of viral replication, some subgroups, such as H5/N8, H7/N1, and H9/N2, display distinct dynamics, possibly reflecting variations in immune response or viral replication capacity.

Transmission odds ratios were estimated for four subgroups: H1/N1, H2/N3, H7/N9, and H9/N2. Although no statistical differences were found between the confidence intervals, H1/N1 exhibited the highest transmission probability, while H7/N9 the lowest one (Table 2). From those highly pathogenic IAV, only ferrets contaminated with the subgroup H7/N9 showed transmission events. On the other hand, daily recovery rates were quite similar across all the evaluated subgroups, ranging from 0.11 to 0.15. The only exception was H5/N8, which had a higher recovery rate of 0.23. However, an overlap of the confidence was also observed. It was only possible to estimate daily mortality rates for five subgroups: H5/N1, H5/N6, H7/N7, H7/N8, H7/N9. The higher mortality rate 0.55 (95% CI 0.42 - 0.68) was found in the H5/N6 subgroup, and the lowest one in the H7/N8 subgroup.

**Table 2.** Estimates of population traits for differents IAVsubgroups.

| HA/NA* | Estimated max viral titer (95% IC) | Transmission rate (95% IC) | Daily recovery rate (day<sup>-1</sup>) (95% IC) | Daily mortality rate (day<sup>-1</sup>) (95% IC) |
| --- | --- | --- | --- | --- |
| H1/N1 | 26.5 (25.6-27.5) | 2.10 (1.25-4.34) | 0.13 (0.12-0.15) | - |
| H2/N2 | 27.4 (24.6-30.1) | - | 0.14 (0.10-0.19) | - |
| H2/N3 | 28.8 (25.9-31.7) | 2.04 (1.24-4.04) | 0.12 (0.08-0.16) | - |
| H3/N2 | 26.2 (23.7-28.8) | - | 0.13 (0.09-0.18) | - |
| H5/N1 | 22.4 (20.4-24.4) | - | 0.13 (0.10-0.17) | 0.41 (0.38-0.45) |
| H5/N2 | 20.3 (16.7-23.8) | - | 0.15 (0.10-0.25) | - |
| H5/N6 | 18.8 (15.6-21.9) | - | 0.14 (0.09-0.22) | 0.55 (0.42-0.68) |
| H5/N8 | 16.9 (12.4-21.4) | - | 0.23 (0.12-0.47) | - |
| H7/N1 | 27.6 (25.1-29.8) | - | 0.12 (0.07-0.20) | - |
| H7/N2 | 27.4 (25.1-29.8) | - | 0.13 (0.10-0.18) | - |
| H7/N3 | 28.9 (26.1-31.8) | - | 0.11 (0.08-0.16) | - |
| H7/N7 | 28.6 (26.2-31.1) | - | 0.12 (0.08-0.16) | 0.29 (0.21-0.36) |
| H7/N8 | 22.9 (19.3-26.4) | - | 0.13 (0.08-0.20) | 0.16 (0.04-0.28) |
| H7/N9 | 28.0 (25.6-30.4) | 1.90 (1.20-3.54) | 0.12 (0.08-0.16) | 0.33 (0.26-0.40) |
| H9/N2 | 33.4 (30.9-36.0) | 1.96 (1.27-3.57) | 0.11 (0.08-0.15) | - |

\*Hemagglutinin/Neuraminidase subgroups; A dash means “not available”.

The geometric mean of cumulative viral titer reached values close to 30 (log10 titer/mL) for subgroups H7/N9 and H7/N7 (Figure 2). The cumulative viral titer for H7/N8 was smaller than both H7/N9 and H7/N7 and fatality was smaller. By contrast, for subgroups H5/N1 and H5/N6 cumulative viral titer was also smaller, however fatality rates were higher. The intervals for the cumulative title have overlaps. The subgroups that did not have any deaths were re-grouped together and also reached values over 25 (log10 titer/mL), even though these many subgroups are quite diverse. In particular, the fatality rates were smaller for subgroups with hemagglutinin HA5 (H5/N2 and H5/N8) and higher for the subgroup H9/N2. Therefore, cumulative titer values were high for fatality rates in intermediate values between 20-40%.

**Figure 2.**
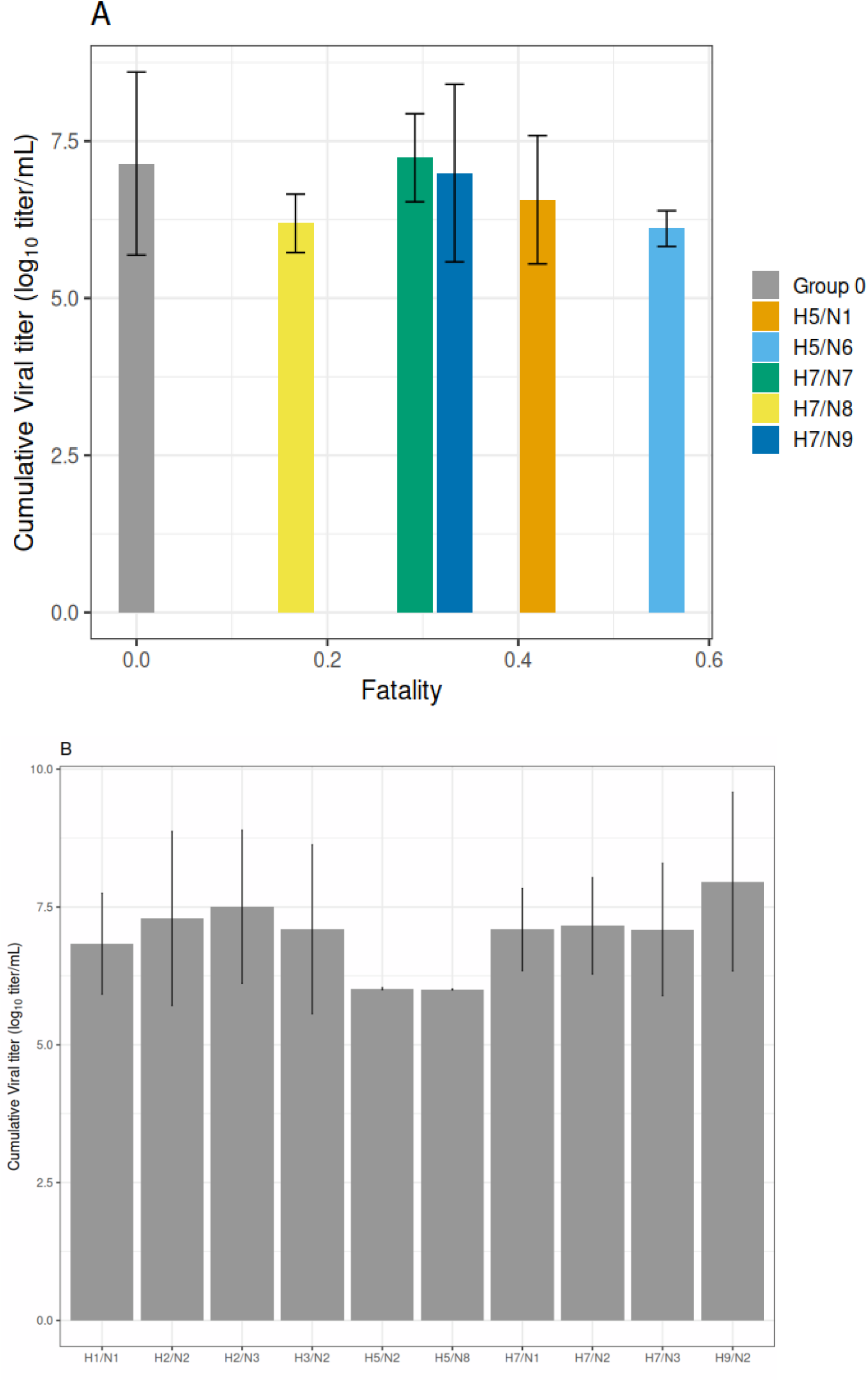
Relationship between cumulative viral titer and fatality for each IAV subgroup (A). Cumulative virtual titer (B) within the subgroups that had fatality rate equal to 0 (Group 0). Bars show values by geometric mean and lines the confidence intervals.

## Discussion

The tradeoff theory in the evolution of virulence posits that virulence evolves toward an intermediate level. Evidence supporting the tradeoff theory is challenging to observe in practice (9), often due to limitations in time and sampling. In this study, the evaluation of cumulative viral titers following IAV infection in ferrets suggests that subgroups infected with IAV strains carrying HA subtype HA 7 and NA subtypes NA 7 and NA 9 were able to accumulate higher viral titers throughout the infection period. Animals in subgroup H5/N6 had the highest fatality rate, along with pronounced weight loss and temperature increases. More importantly, the subgroup H5/N6 did not reach high cumulative viral titers. However, when examining the cumulative viral titer using geometric means, levels were slightly lower than those observed in ferrets infected with subgroups H7/N7, H7/N9, and H5/N1, though there is some uncertainty due to sample size constraints. Those with H7/N3 and H7/N9 showed both high weight loss and increased temperature. For subgroup H7/N9, the fatality rate exceeded 30%, whereas no fatalities were observed in the H7/N3 subgroup. Additional measures, such as elevated temperature, were also higher in these subgroups, translating into more symptomatic events with higher cumulative titers, as expected. When cumulative viral titer is considered a proxy for the reproduction number (i.e., the potential secondary infection rate), the findings suggest an intermediate level as given in the hypothesis.

The cumulative viral titer counted from infection day to the lethal day, if any, gives a total measure of the transmissibility during the course of infection. The expectation is that individuals with higher cumulative viral titers will transmit to more individuals that are eventually in contact. Therefore, these values can be counted as proxies to reproduction numbers. The results suggest that higher fatality rates impose restriction on the number of potential secondary cases. By contrast, subgroups with no fatalities also reached high values of cumulative titer, comparable to subgroups H7/N9 and H7/N7. From this perspective, lower fatality rates did not cause a significantly higher potential for secondary cases, although uncertainty levels make confirmation difficult.

Daily evaluations of viral titers indicate that, in some subgroups, high viral titers were sustained for approximately five days post-infection. For most subgroups, viral titers declined substantially after day seven, and cumulative levels were higher when IAV titers were sustained over a longer period. Subgroups such as those infected with subtype H5 that did not initially show high viral titers also did not achieve high cumulative titers. Similarly, subgroups with faster viral decline, such as those with subtypes H1 and H2, did not reach high cumulative titers. Notably, subgroup H9/N2 had the highest mean titer, yet no fatalities were observed in this subgroup.

Mortality rates were also examined and ranking among the subgroups is similar to the ranking according to fatality as expected. For H5/N6, in particular, the confidence levels indicate that mortality is significantly higher for animals infected with this subgroup, also placing it in the end spectrum of advantage for persistence.

The theory behind the tradeoff hypothesis (4,10,11) can be explained via a mathematical model in which individuals go on stages of infection, recovery, and mortality. The transmissibility is given as a function of the virulence, given by mortality, however equal increases of mortality lead to diminished increases in transmissibility as virulence increases. This theoretical explanation has biological plausibility and leads to an optimal level of transmissibility that can be evaluated at least in theory via the reproduction number.

Fatality by AIV is not only caused by the direct effects of infection, such as the destruction of infected epithelial cells. It is also linked to the cytokine storm, which correlates directly with viral titer levels. In humans, fatality associated with the H5/N1 subgroup is partly due to its ability to infect various tissue types, leading to a more intense viral titer and cytokine storm (16,17). However, this subgroup had the second-highest fatality despite one of the lowest cumulative viral titers among the subgroups (Figure 2A). A similar pattern was observed in the H5/N6 subgroup, which had the highest fatality, yet also showed the lowest cumulative viral titers, suggesting that possibly these subgroups had not evolved to an optimal level of virulence yet. Another evidence is that none of the ferrets inoculated with these subgroups had directly transmitted to a naive one, which is in accordance with these highly pathogenic avian influenza (HPAI) (18,19). This type of transmission only occurred with the H5/N1 subgroup after genetic modifications and several passages in ferrets, which, when inoculated with these modified H5/N1 strains, exhibited significantly lower lethality (20).

On other hand, subgroups related to subtype HA 7 like H7/N1, H7/N3, and H7/N7 had been responsible for more than 100 million poultry deaths in this century in 4 continents and 10 countries (21). In the results, ferrets inoculated with subgroups H7/N7 and H7/N9 had the highest cumulative titer and demonstrated a lower fatality, which can possibly be a sign of a trade-off hypothesis. However, transmission of this subgroup was not seen in the original sample, but only 25% percent of ferrets had these results (Table 1). Direct transmission occurs in another animal model after mutations related to the polymerase subunit of these viruses (22). In this way, HA H7, especially H7/N7 is a possible figure as one of the subgroups more related to optimal virulence and transmission between their original poultry hosts.

There is evidence of the tradeoff hypothesis with different pathogens. Most notably, the frequency of myxoma virus has been shown to evolve to groups with intermediate virulence grade in Australia in the 1950s and 1960s (12,13). Also in Britain, where an intermediate-virulence strain of myxoma tends to be more common in two samples collected six years apart (14). Works that covered cohorts of individuals with HIV indicate long periods of virus incubation. Fraser *et al*., 2007 (15) examined the distribution of set-point viral loads, where Individuals with intermediate levels of set-point viral loads had higher transmission potential. In particular, Fraser *et al*., 2007 (15) also examined viral loads and obtained an estimated number of secondary infections from the studied cohort.

This study was constrained in the analysis due to the number of samples. Transmissibility could be measured via an indicator contained in the original dataset about transmission via respiratory droplets. Transmissibility as stated in theoretical works could vary according to virulence. However, the results were limited due to the evaluation of this transmission in a much smaller subsample of animals in the studied subgroups. The original study was not initially designed to evaluate a comparison of virulence levels across the different subtypes. However, the analysis considered more than 350 individuals which permitted obtaining significant results. Additional studies of the influenza virus in animal models are recommended.

Knowledge of infectious diseases in animals provides evidence valuable for animal models and from a One Health perspective. When spillover events occur, transferring pathogens from animals to human populations, an integrated health approach has increasingly guided advancements across all related areas. The present study suggests significant virulence of IAV in ferrets with potential for transmissibility.

## Acknowledgements

DAMV is grateful for the National Council for Scientific and Technological Development (CNPq/Brazil, Ref. 312282/2022-2), Fundação Carlos Chagas Filho de Amparo à Pesquisa do Estado do Rio de Janeiro (Ref. E-26/204.108/2024), and CAPES (Service Code 001). ASG and LGM receive PhD scholarships from National Council for the Improvement of Higher Education Personnel (CAPES/Brazil).

## Author contributions

**Ayrton S. Gouveia**: Writing - original draft, Writing – review and editing, Data curation, Formal analysis, Investigation, Methodology, Software, Visualization; **Leonardo G. Mata**: Writing - original draft, Writing – review and editing, Data curation, Formal analysis, Investigation, Methodology, Software, Visualization; **Daniel A. M. Villela**: Writing - original draft , Writing – review and editing, Conceptualization, Data curation, Investigation, Methodology, Project administration, Software, Supervision, Validation, Visualization

